# Curcumin induces mild anemia in a DSS-induced colitis mouse model maintained on an iron-sufficient diet

**DOI:** 10.1101/479428

**Authors:** Macha Samba-Mondonga, Marco Constante, Gabriela Fragoso, Annie Calvé, Manuela M. Santos

## Abstract

Anemia is frequently encountered in patients with inflammatory bowel disease (IBD), decreasing the quality of life and significantly worsening the prognosis of the disease. The pathogenesis of anemia in IBD is multifactorial and results mainly from intestinal blood loss in inflamed mucosa and impaired dietary iron absorption. Multiple studies have proposed the use of the polyphenolic compound curcumin to counteract IBD pathogenesis since it has significant preventive and therapeutic properties as an anti-inflammatory agent and very low toxicity, even at high dosages. However, curcumin has been shown to possess properties consistent with those of an iron-chelator, such as the ability to modulate proteins of iron metabolism and decrease spleen and liver iron content. Thus, this property may further contribute to the development and severity of anemia of inflammation and iron deficiency in IBD. Herein, we evaluate the effects of curcumin on systemic iron balance in the dextran sodium sulfate (DSS) model of colitis in C57Bl/6 and BALB/c mouse strains that were fed an iron-sufficient diet. In these conditions, curcumin supplementation caused mild anemia, lowered iron stores, worsened colitis and significantly decreased overall survival, independent of the mouse strain. These findings suggest that curcumin usage as an anti-inflammatory supplement should be accompanied by monitoring of erythroid parameters to avoid exacerbation of iron deficiency anemia in IBD.

**Core tip:** Our study shows that curcumin supplementation of an iron-sufficient diet causes mild anemia, depletes iron stores, worsens colitis and decreases survival, independent of the mouse strain. These findings demonstrate that curcumin may have a potential adverse activity in chronic inflammatory diseases, such as IBD, since the iron chelating properties of curcumin may play a role in iron deficiency and the severity of colitis.

## Introduction

For patients with inflammatory bowel disease (IBD), anemia is one of the major causes of hospitalization (27, 32) and has a debilitating effect on the quality of life (QoL) (47, 54), increasing disease morbidity and tightly associating the disease with mortality (12). IBD is an inflammatory disease consisting of a group of gastrointestinal tract disorders, namely ulcerative colitis and Crohn's disease, which are characterized by blood loss from the intestinal mucosa and reduced iron absorption. Up to two-thirds of patients with IBD develop anemia, with the most common types being iron-deficiency anemia and anemia of chronic disease, which often overlap (27, 40, 46).

IBD pathophysiology includes the activation of inflammatory cytokines such as TNF-α (34, 42). Therefore, many studies on IBD therapies have focused on anti-inflammatory treatments or natural compounds such as curcumin that have anti-inflammatory properties to mitigate the disease (5, 13).

Curcumin, the yellow pigment obtained from the rhizome of *Curcuma longa* (turmeric), is commonly used as a spice and food-coloring agent (14). Curcumin features complex and multifactorial mechanisms of action that have demonstrated a variety of therapeutic properties, including those described as anti-oxidant, anti-infection, anti-tumor, and anti-inflammatory (49). Moreover, curcumin use has no major side effects and has low toxicity at high dosages (8-12 g/day) (8). Anti-cancer activities of curcumin are mediated by a variety of biological pathways in mutagenesis, oncogene expression, cell cycle regulation, apoptosis, tumorigenesis, and metastasis. Additionally, the effects of curcumin as an anti-inflammatory agent have been previously associated with the regulation of different inflammatory cytokines (extracellular matrix metalloproteinase inducer, EMMPRIN; matrix metalloproteinase-9, MMP-9; IL-1β; and mitogen-activated protein kinase, MAPK) (6, 39, 51). Recently, Kong *et al.* demonstrated curcumin’s mechanism of action in macrophages during the inflammasome response (26). Furthermore, different studies have highlighted curcumin’s properties to inhibit the toll-like receptor 4/myeloid differential factor 88 (TLR4/MyD88) pathway via the repression of TLR4 homodimerization and the subsequent decrease of MyD88 expression (58, 59). TLR4 has a critical role in the inflammatory response inducing nuclear factor-kappa B (NF-κB) expression via protein adaptor MyD88 stimulation.

Curcumin has also been shown to have anticoagulant and antiplatelet activities (24), which may sustain or prolong active bleeding (25) and has been proposed to have the properties of an iron chelator (4, 22, 23). Consistent with its iron chelating properties, curcumin has been reported to reduce spleen and liver iron stores in mice (9, 23). Moreover, curcumin has been also shown to affect hepcidin expression (23), the main regulator of iron homeostasis (18). Hepcidin, encoded by the *HAMP* gene, controls the levels of intestinal iron absorption and plays a major role in regulating iron release from macrophages. These cells are responsible for iron recycling and, in inflammatory settings, will accumulate iron at high hepcidin levels (53).

The potentially detrimental effects of curcumin on iron homeostasis in the inflammatory context could exacerbate anemia and iron deficiency; however, this aspect has been often ignored in studies of gastrointestinal disorders and IBD mouse models, which exhibit marginal or depleted iron stores. Herein, we investigated the effects of curcumin on the dextran sodium sulfate (DSS)-induced colitis mouse model in the context of an iron-sufficient diet.

## Materials and Methods

### Animals

All procedures were performed in accordance with the Canadian Council of Animal Care guidelines after approval by the Institutional Animal Care Committee of the Centre de recherche du Centre Hospitalier de l'Université de Montréal (CRCHUM). Female C57Bl/6 and BALB/c mice aged 8 weeks old were purchased from Charles River Laboratories (Wilmington, CA, USA). Animals were maintained under standard 12:12 light/dark conditions at the CRCHUM.

### Animal treatments, diet, and induction of colitis

Two weeks before DSS treatment, 8 weeks old female C57BL/6 or BALB/c mice received *ad libitum* a control diet containing 50 mg/kg of iron in the form of iron sulfate (Teklad TD.120515; Envigo, IN, United States) with or without 2% (wt/wt) curcumin supplementation (Teklad TD.140182). Colitis was induced by the administration of DSS (molecular weight 40000; TdB Consultancy, Uppsala, Sweden) at 1-2% w/v in drinking water for 5 days, followed by 7 days of rest for 3-4 cycles of DSS (10).

### Erythroid parameters and serum iron

Red blood cell, hemoglobin, hematocrit, and mean corpuscular volume levels were measured with an automated cell counter calibrated for murine samples (ABC vet counter; ABX Hématologie, Montpellier, France). Serum iron was measured by a colorimetric assay with the Kodak Ektachem DT60 system (Johnson & Johnson, Ortho-Clinical Diagnostics, Mississauga, ON, Canada).

### Histology

Histological scoring was assessed on colon samples from each mouse. The samples were fixed in 10% neutral buffered formalin (Chapter Chemicals, Montreal, QC, Canada), cut, fixed and stained with hematoxylin and eosin. All histological evaluations were assessed in a blinded fashion. Histological scoring was calculated as follows: presence of occasional inflammatory cells in the lamina propria (assigned a value of 0); increased numbers of inflammatory cells in the lamina propria (value of 1); confluence of inflammatory cells, extending into the submucosa (value of 2); and transmural extension of the infiltrate (value of 3). For tissue damage score: no mucosal damage (value of 0); lymphoepithelial lesions (value of 1); surface mucosal erosion or focal ulceration (value of 2); and extensive mucosal damage and extension into a deeper structure (value of 3) (21).

### Disease activity index

The disease activity index was scored accordingly to previous studies (11, 15). The index consists of the sum of all scores attributed to weight loss (0, none; 1, 1%–5%; 2, 5%– 10%; 3, 10%–20%; 4, >20%), stool consistency (0, normal; 2, soft; 4, diarrhea) and fecal blood (0, negative; 2, blood in the stool; 4, gross bleeding), divided by 3.

### SDS-PAGE and western blot analysis

Protein concentrations were measured using a Pierce BCA protein assay kit (Thermo Fisher Scientific, Rockford, IL, USA). Same concentrations of protein from colon or liver extracts were loaded to perform electrophoresis. More precisely, equivalent amounts of proteins were boiled in loading buffer containing 4% SDS, 20% glycerol, and bromophenol blue for 5 minutes. Proteins were resolved on 10% SDS-PAGE gels and then were transferred onto nitrocellulose membranes (GE Healthcare; Amersham Biosciences, Baie d’Urfé, QC, Canada). The membranes were blocked with 5% non-fat dry milk solution and incubated with antibodies against lipocalin 2 (R&D Systems, Minneapolis, MN, USA), MyD88 (Cell Signaling, Danvers, MA, USA) and β-actin (Abcam, Cambridge, MA, USA). To detect the formation of immunocomplexes, peroxidase-conjugated anti-goat IgG (Santa Cruz, Dallas, TX, USA) and anti-mouse IgG (R&D Systems, Minneapolis, MN, USA) were used as secondary antibodies. Staining intensity was developed with an Amersham enhanced chemiluminescence system (GE Healthcare, Amersham Biosciences, Baie d’Urfé, QC, Canada).

### Quantitative RT-PCR

Total RNA from tissue samples was isolated by phenol chloroform using the TRIzol reagent (Invitrogen, Burlington, ON, Canada) as recommended by the manufacturer, and reverse transcription was performed with an Omniscript RT-PCR system (Qiagen, Mississauga, ON, Canada). The mRNA levels of selected genes were measured by real-time PCR with a Rotor-Gene 3000 real-time DNA detection system (Montreal Biotech, Kirkland, QC, Canada) and QuantiTect SYBR Green I PCR kits (Qiagen, Mississauga, ON, Canada) as previously described (35). Expression levels were normalized to the housekeeping gene β-actin. The following primers were used: *Hamp* (F) CCTATCTCCATCAACAGATG; *Hamp* (R) AACAGATACCACACTGGGAA; β-Actin (F) TGTTACCAACTGGGACGACA; β-Actin (R) GGTGTTGAAGGTCTCAAA.

### Measurement of iron in the liver and the spleen

Non-heme iron concentrations were assessed by acid digestion of liver and spleen tissue samples (56), followed by measurement by colorimetry using the ferrozin reagent and measuring absorption at 560 nm (17).

### Statistical analysis

All statistics were calculated using Prism GraphPad (GraphPad, San Diego, CA) with a pre-specified significant *P*-value of 0.05. Student’s *t*-test (two-tailed) was used for comparisons between two groups, and multiple comparisons were evaluated by one-way analysis of variance (ANOVA), followed by the Bonferroni multiple comparison test.

## Results

### Curcumin supplementation of an iron-sufficient diet causes mild anemia in a DSS-induced colitis mouse model

We investigated the potential contribution of curcumin in the systemic iron balance in an IBD setting using the DSS-induced colitis mouse model. To evaluate such effects, colitis was induced by administration of DSS in C57Bl/6 mice with or without 2% curcumin supplementation of an iron-sufficient diet (50 mg/kg chow). Repeated cycles of DSS administration in the drinking water of mice were intercalated with resting periods of 7 days with untreated? water to model the chronic pattern of IBD (45, 55).

We found that erythroid parameters including red blood cells, hemoglobin, mean corpuscular volume and hematocrit values were significantly lower in DSS-treated mice on the curcumin-supplemented diet (Curcumin-DSS), compared with mice treated with DSS without curcumin supplementation (Ctrl-DSS) **(figure 1A-D)**. In control conditions with non-colitic mice (Ctrl and Curcumin, without DSS), only the mean corpuscular volume was lowered by curcumin supplementation, with all other erythroid parameters remaining similar in both groups. The lowest values for serum iron levels were found in the Curcumin-DSS group, with levels significantly lower compared to non-colitic Ctrl mice **(figure 1E)**.

**Figure 1.**
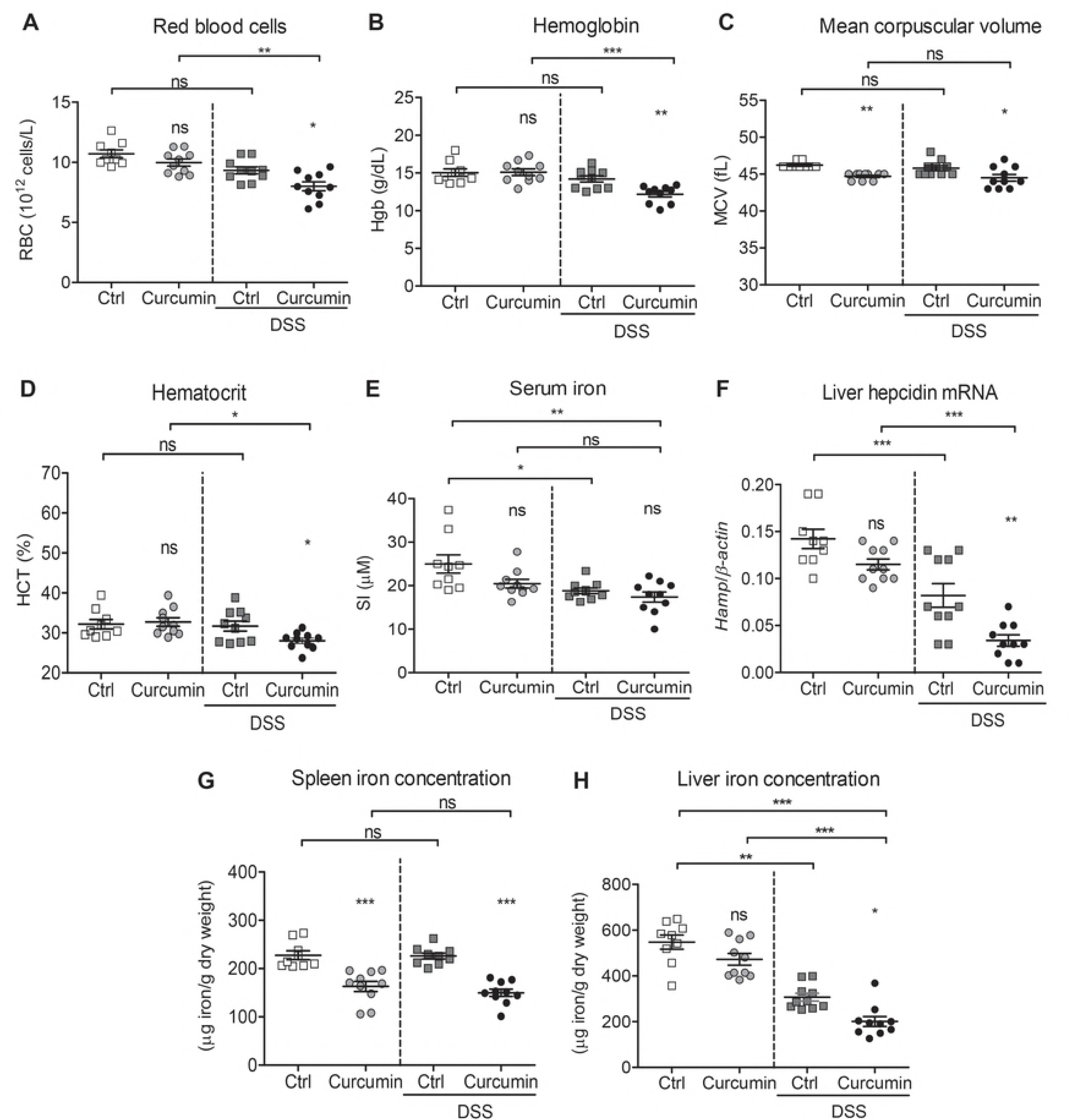
Curcumin supplementation of an iron-sufficient diet causes mild anemia in a DSS-mouse model. C57BL/6 mice were fed an iron-sufficient diet (50 mg/kg chow; Ctrl) or an iron-sufficient diet supplemented with 2% curcumin (Curcumin). For dextran sodium sulfate (DSS) treatment, mice were fed an iron-sufficient diet with or without curcumin, starting 2 weeks before administration of DSS. (A-D) Erythroid parameters: red blood cells, hemoglobin, mean corpuscular volume, and hematocrit. (E) Serum iron levels. (F) Liver hepcidin (*Hamp)* mRNA expression against housekeeping β-actin mRNA. (G-H) Iron content in spleen (G) and liver (H). Results are representative of a minimum of three independent experiments; n = 9-10 mice per group. Statistical analysis was performed with one-way ANOVA. *\*P* < 0.05, *\*\*P* < 0.01, *\*\*\*P* < 0.001, and ns = not significant between Curcumin to Ctrl groups and when indicated between non-DSS and DSS groups.

Next, we measured liver hepcidin mRNA expression *(Hamp*), the major regulator of iron homeostasis (18). As shown in **figure 1F,** Curcumin-DSS mice had the lowest hepcidin expression of all groups, and the values were also significantly lower when compared to Ctrl-DSS mice that did not received curcumin supplementation.

We then analyzed the iron concentration in the spleen and liver and found that splenic iron concentrations in mice fed with 2% curcumin with or without DSS treatment were significantly lower compared with their respective controls (Ctrl and Ctrl-DSS; **figure 1G**). In regards to liver concentration, both groups treated with DSS had lower liver iron content compared to non-DSS mice, indicating that DSS treatment lowers liver iron concentrations **(figure 1H)**. The lowest liver iron content was found in the Curcumin-DSS group when compared to all other treatment groups.

Taken together, these findings indicate that mice fed an iron-sufficient diet supplemented with curcumin develop mild anemia accompanied by marked lower iron levels in the spleen and liver in the DSS-mouse model. In contrast, mice treated with DSS alone do not develop anemia despite a modest lowering of serum and liver iron concentrations.

### Curcumin supplementation aggravates colitis in the DSS-mouse model

We evaluated the effect of curcumin supplementation on colitis severity. Curcumin supplementation in DSS-treated mice contributed to a greater weight loss compared to Ctrl-DSS mice **(figure 2A)**. This result was in line with survival **(figure 2B)**, where both Ctrl-DSS and Curcumin-DSS groups demonstrated mortality. However, only Curcumin-DSS mice showed a steady decrease in survival reaching 60% compared to 90% in the Ctrl-DSS group at the end of the experiment.

**Figure 2.**
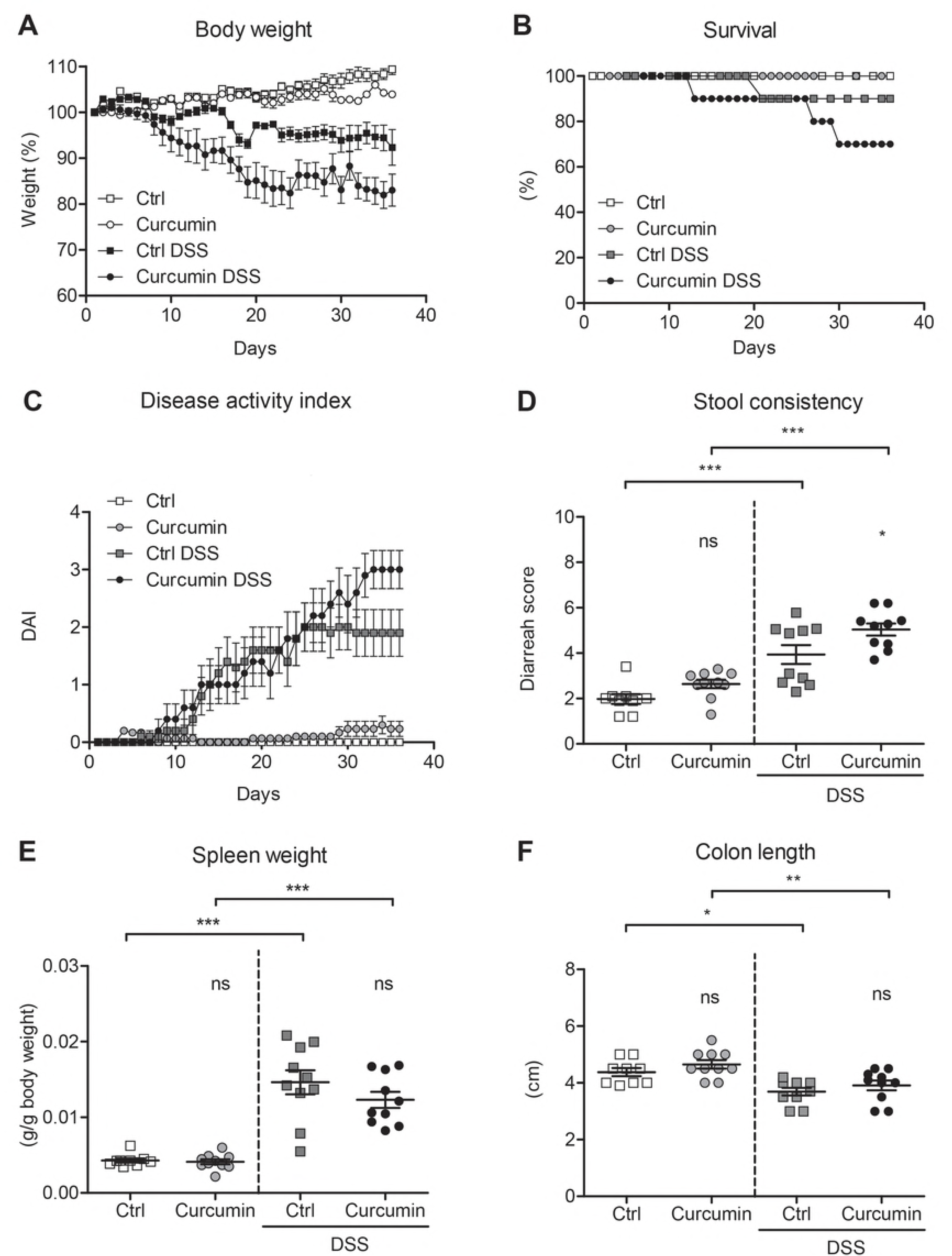
Curcumin supplementation of an iron-sufficient diet aggravates colitis in a DSS-mouse model. C57BL/6 mice were fed an iron-sufficient diet (Ctrl) or iron-sufficient diet supplemented with curcumin (Curcumin). For dextran sodium sulfate (DSS) treatment, mice were fed an iron-sufficient diet with or without curcumin, starting 2 weeks before administration of DSS. (A) Body weight. (B) Survival. (C) Disease activity index (DAI). (D) Stool consistency. (E) Spleen weight. (F) Colon length. Results are representative of a minimum of three independent experiments; n = 9-10 mice per group. Statistical analysis was performed with one-way ANOVA. *\*P* < 0.05, *\*\*P* < 0.01, *\*\*\*P* < 0.001, and ns = not significant between Curcumin to Ctrl groups and when indicated between non-DSS and DSS groups.

The disease activity index, consisting of the sum of the scores attributed to body weight loss, stool consistency and observance of blood in feces divided by three, was measured according to previous studies (11, 15). We found that Curcumin-DSS mice had an increased disease activity index score compared to Ctrl-DSS mice **(figure 2C)** after 30 days of treatment. The stool consistency score, highest for diarrhea, reflected the same trend and showed a significant increase in Curcumin-DSS mice compared to Ctrl-DSS **(figure 2D)**.

Spleen weight was significantly higher in both groups of mice treated with DSS (Ctrl-DSS and Curcumin-DSS) than non-colitic mice **(figure 2E)**, with no significant differences found between Ctrl-DSS and Curcumin-DSS. We additionally analyzed the colon length as a marker of disease severity and found that it was significantly shorter in DSS-treated mice (Ctrl-DSS and Curcumin-DSS) than in non-colitic mice, but curcumin treatment did not did not appear to enhance the shortening of the colon **(figure 2F)**.

Overall, these results show that, in our experimental setting, curcumin supplementation decreases survival and is associated with higher scores of disease activity in DSS-induced colitis.

### Curcumin supplementation in DSS-treated mice enhances inflammation in the colon

We assessed protein levels of lipocalin 2 in the colon as it is highly expressed in response to injury and inflammation and is used as a biomarker of intestinal inflammation (7). Our results showed that curcumin supplementation of DSS-treated mice was associated with enhanced lipocalin 2 expression **(figure 3A-B)** compared to Ctrl-DSS mice, indicating heightened inflammation.

**Figure 3.**
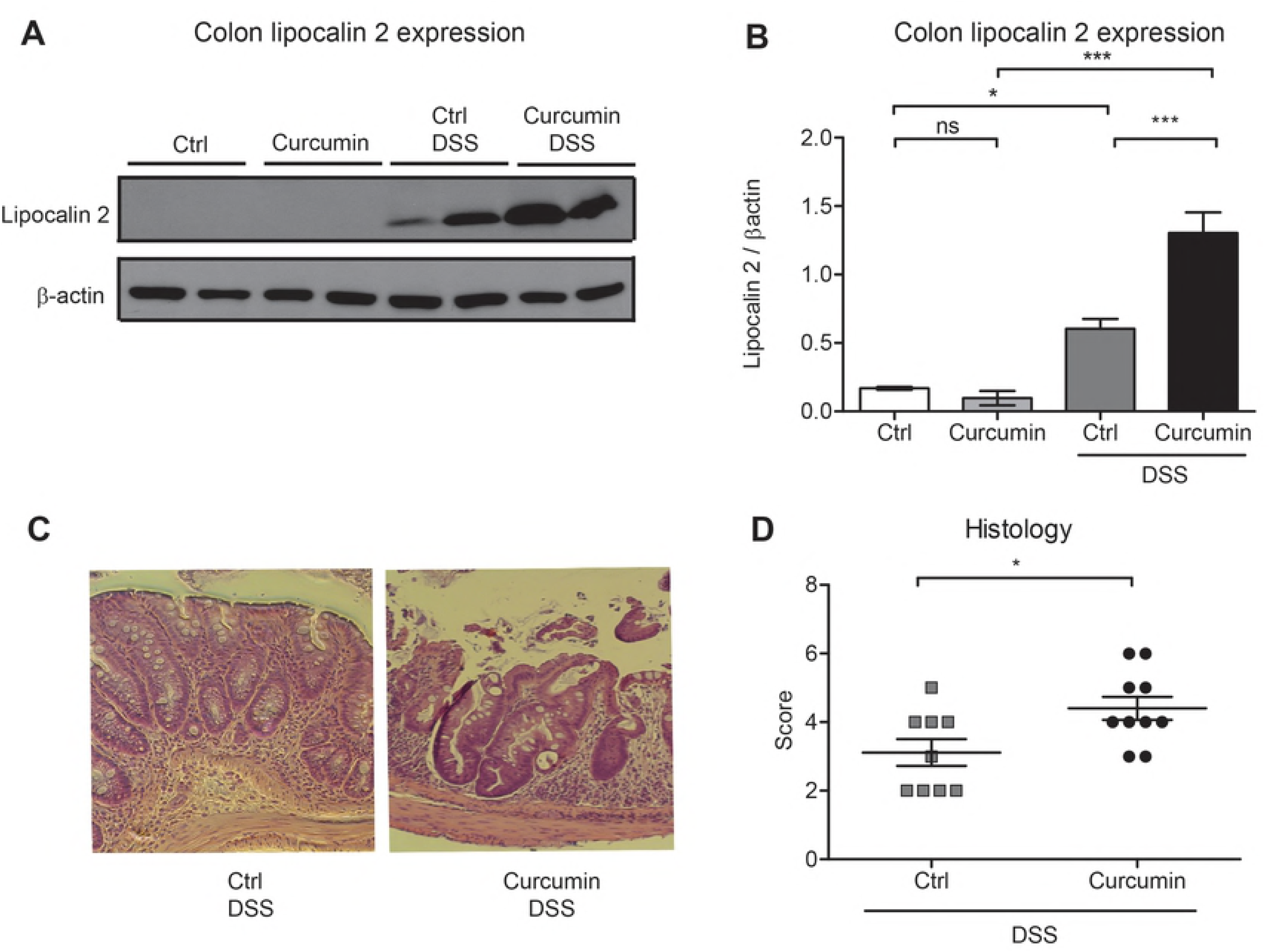
Curcumin supplementation in DSS-treated mice enhances inflammation in the colon. (A) Representative western blot of colon protein extracts probed with antibodies against lipocalin 2 and β-actin. (B) Graphic depicting densitometric quantification of western blots from three independent experiments. Data are presented as mean ± SEM. (C) Representative hematoxylin and eosin staining of mouse colon histological sections. (D) Graphic depicting quantification of colonic histology scores. Statistical analysis was performed by two-tailed Student’s *t*-test. *\*P* < 0.05, *\*\*P* < 0.01, *\*\*\*P* < 0.001; ns = not significant; n = 8 mice per group.

We then assessed the severity of DSS-induced colitis using histology scoring (see materials and methods). We found that cellular infiltration and tissue damage followed by epithelial destruction were more severe in mice supplemented with curcumin (Curcumin-DSS) compared to mice treated with DSS without curcumin supplementation (Ctrl-DSS; **figure 3C-D)**.

We also analyzed curcumin modulation of MyD88 protein expression. MyD88 is the major adaptor protein essential for the inflammatory cytokine activation upon stimulation of almost all the TLRs except for TLR3 (28). As shown in **figure 4A-B**, DSS treatment resulted in MyD88 activation (Ctrl vs. Ctrl-DSS groups). Curcumin supplementation significantly decreased MyD88 expression in colonic tissues in both Curcumin and Curcumin-DSS groups compared to Ctrl and Ctrl-DSS mice. In contrast to the colon, no significant modulation of MyD88 was observed in the liver between the four mouse groups **(figure 4C-D)**.

**Figure 4.**
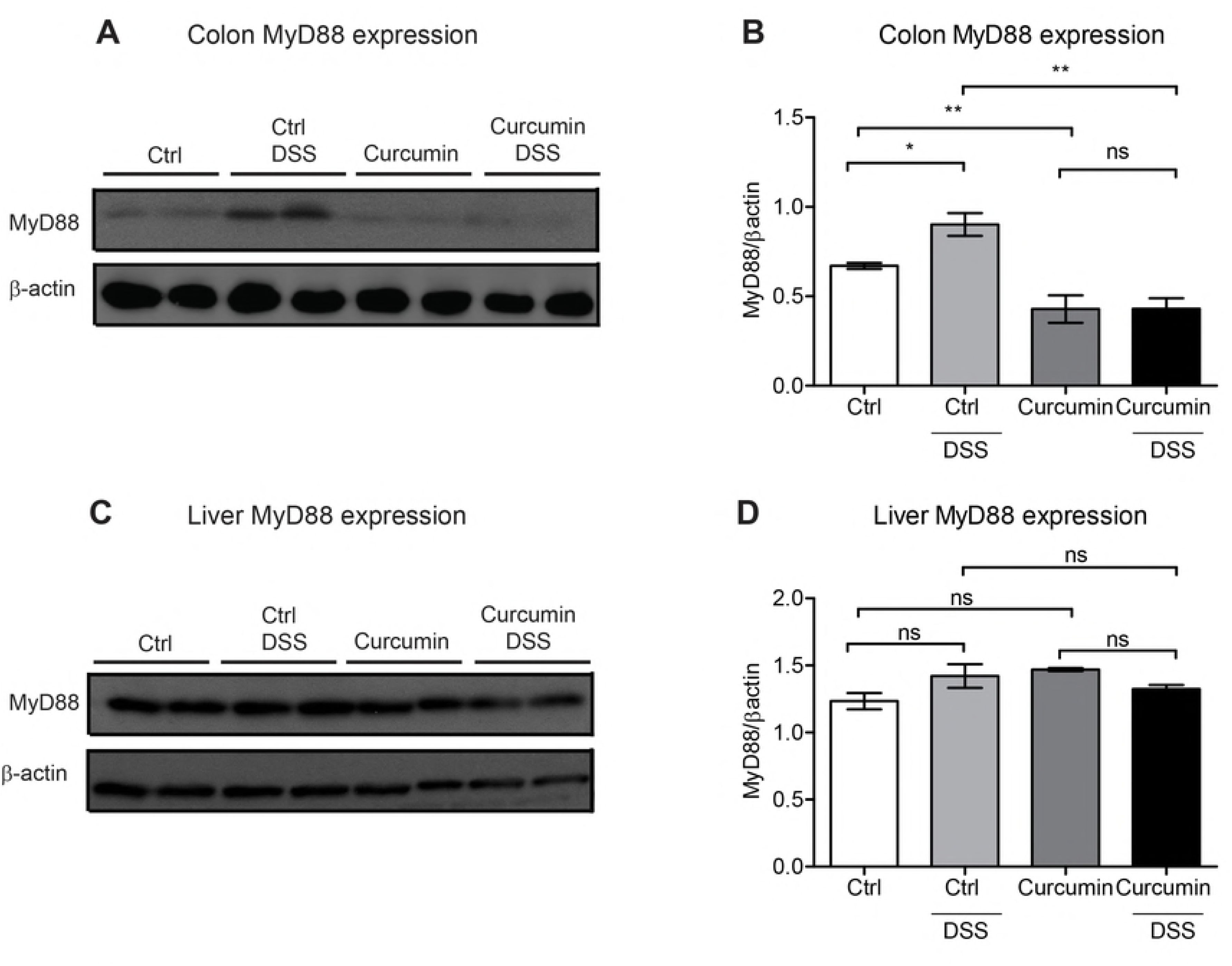
Curcumin decreases MyD88 protein expression in the colon but not in the liver. (A-B) Colonic MyD88 protein expression in C57BL/6 mice treated with or without dextran sodium sulfate (DSS) and fed an iron-sufficient diet (Ctrl) or iron-sufficient diet supplemented with curcumin (Curcumin). (A) Representative western blot of colon protein extracts probed with antibodies against MyD88 and β-actin. (B) Graphic depicting densitometric quantification of western blots from three independent experiments. (C-D) Hepatic MyD88 protein expression. (C) Representative western blot of liver protein extracts probed with antibodies against MyD88 and β-actin. (D) Graphic depicting densitometric quantification of western blots from three independent experiments. Statistical analysis was performed with one-way ANOVA: data in B and C are presented as mean ± SEM. Statistical analysis was performed by two-tailed Student’s *t*-test. *\*P* < 0.05, *\*\*P* < 0.01; ns = not significant; n = 8 mice per group.

These findings show that when mice are fed an iron-sufficient diet, curcumin enhances inflammation and aggravates colitis induced by DSS. In addition, curcumin treatment suppresses MyD88 protein expression in the colon but not in the liver and is independent of DSS induction of colitis.

### Curcumin supplementation of an iron-sufficient diet induces mild anemia independent of mouse strain

To investigate whether the aggravation of colitis in mice by curcumin is dependent on the mouse strain, we assessed the DSS-induced colitis mouse model on BALB/c mice using the same iron-sufficient diets with or without curcumin supplementation at 2% (w/w). We found that BALB/c mice showed, as previously reported (37), higher resistance to the DSS treatment, recovering more readily from the successive DSS-treatment cycles. However, around day 35, body weight loss and survival decreased in Curcumin-DSS mice compared to Ctrl-DSS mice **(figure 5A-B)**. Disease activity index was higher in Curcumin-DSS mice compared to Ctrl-DSS **(figure 5C)**, whereas no significant differences were found in the colon length and spleen weight **(figure 5D-E)**. Lipocalin 2 expression and histology scoring in the distal colon were higher in mice supplemented with curcumin (Curcumin-DSS) than in Ctrl-DSS mice **(figure 5F-I)**, indicating more severe inflammatory cell infiltration and tissue damage in Curcumin-DSS mice.

**Figure 5.**
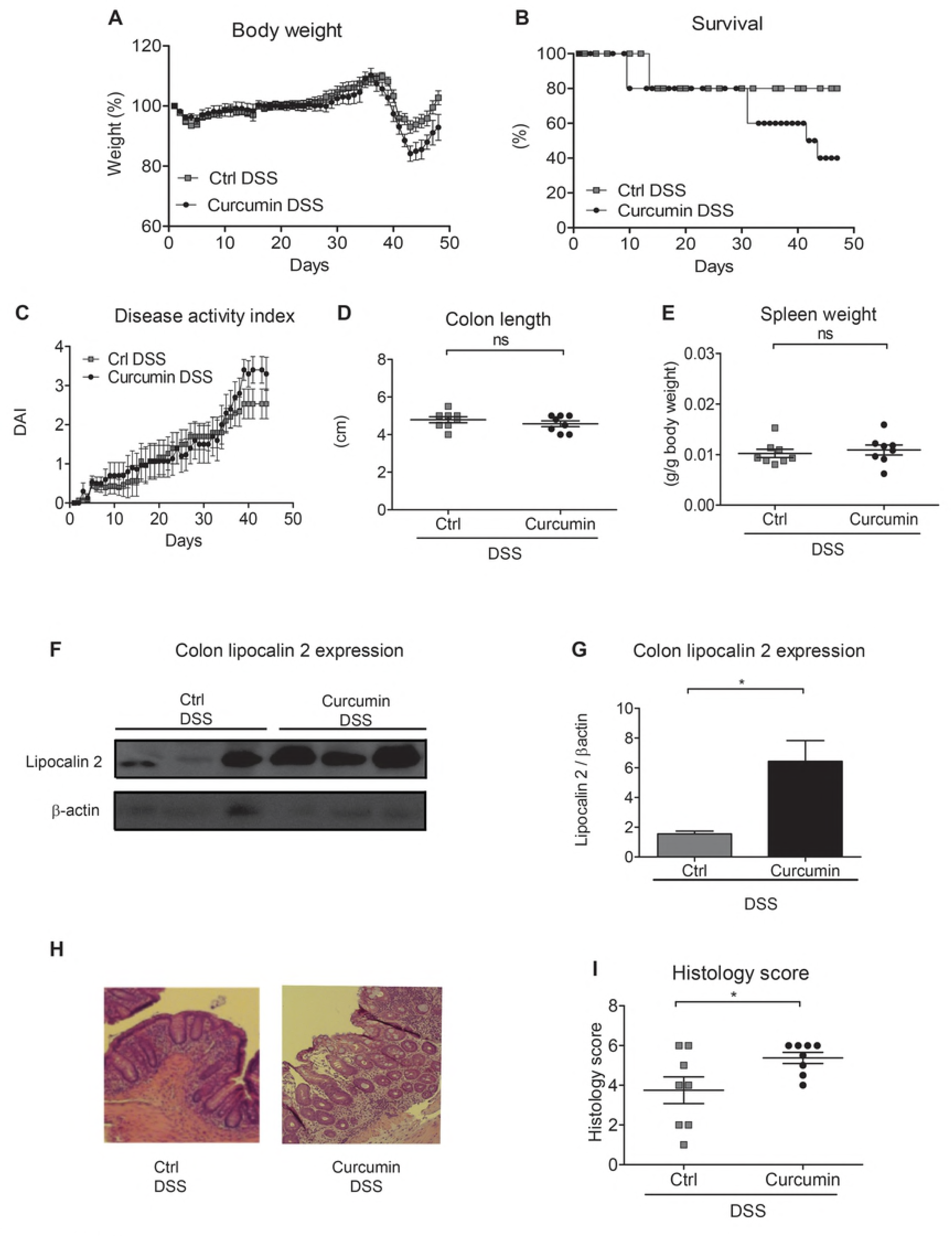
Curcumin supplementation of an iron-sufficient diet in BALB/C mice exacerbates DSS-induced colitis. BALB/c mice were fed an iron-sufficient diet (50 mg/kg chow; Ctrl) or iron-sufficient diet supplemented with 2% curcumin (Curcumin) starting at 2 weeks before administration of dextran sodium sulfate (DSS). (A) Body weight. (B) Survival. (C) Disease activity index (DAI). (D) Colon length. (E) Spleen weight. (F-G) Colonic lipocalin 2 expression. (F) Representative western blot of colonic lipocalin 2 and β-actin expression. (G) Graphic depicting densitometric quantification of western blots from three independent experiments. Data are presented as mean ± SEM. (H) Representative hematoxylin and eosin staining of mouse colon histological section. (I) Graphic depicting quantification of colonic histology scores. Results are representative of a minimum of three independent experiments; n = 8 mice per group. Statistical analysis was performed by two-tailed Student’s *t*-test. *\*P* < 0.05; ns = not significant.

Regarding erythroid parameters, we found that Curcumin-DSS mice had significantly lower red blood cells numbers as well as lower hemoglobin and hematocrit values, while mean corpuscular volume remained unaffected when compared to Ctrl-DSS mice **(figure 6A-D)**. Significant differences were also found in serum iron levels and liver hepcidin expression, with lower values found in Curcumin-DSS than in Ctrl-DSS mice **(figure 6E-F)**. Finally, spleen and liver iron concentrations were significantly lower in Curcumin-DSS mice compared to Ctrl-DSS mice **(figure 6G-H)**.

**Figure 6.**
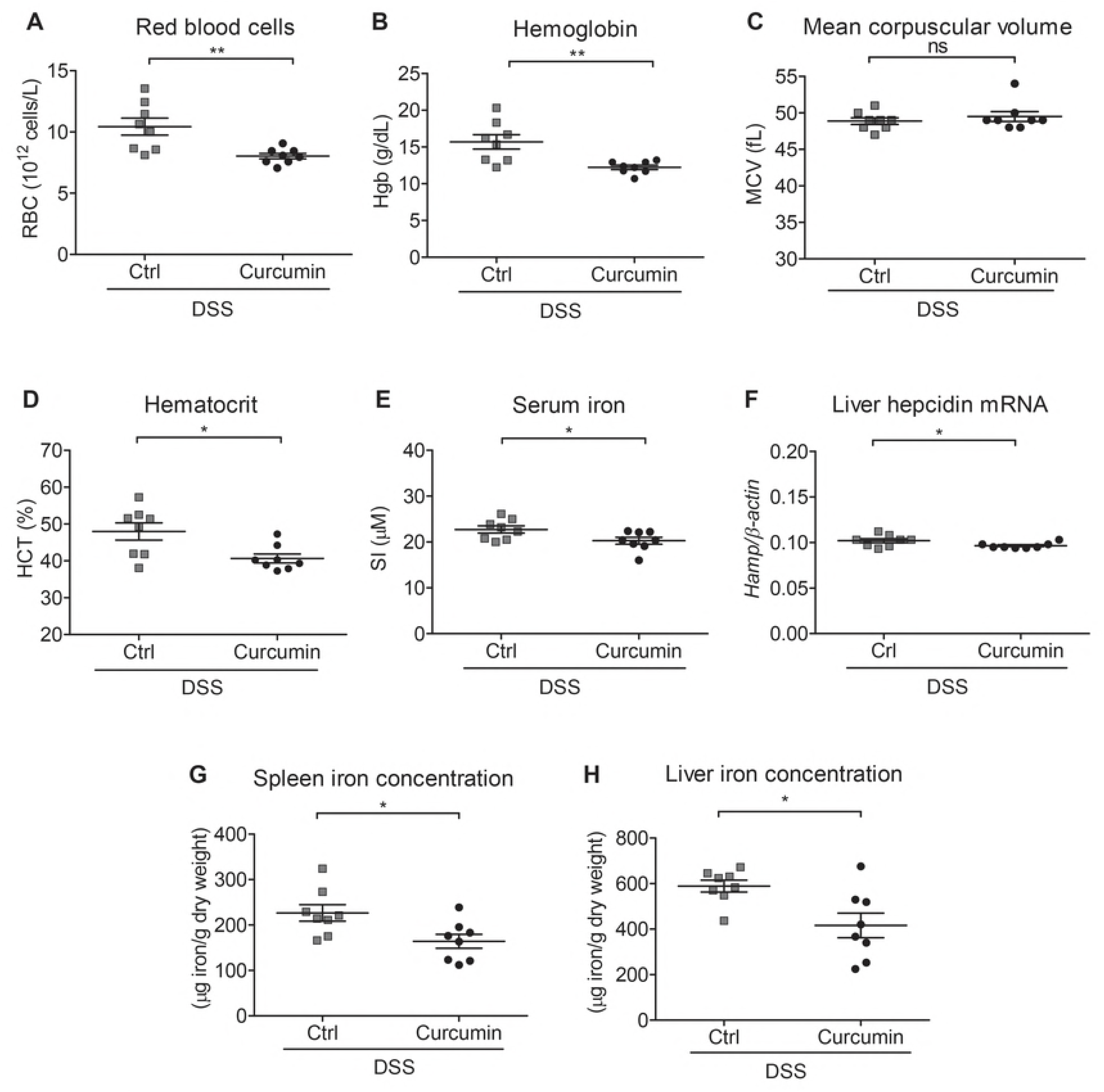
Curcumin supplementation of an iron-sufficient diet induces mild anemia independent of mouse strain. BALB/c mice were fed an iron-sufficient diet (50 mg/kg chow; Ctrl) or iron-sufficient diet supplemented with 2% curcumin (Curcumin) starting at 2 weeks before administration of dextran sodium sulfate (DSS). (A-D) Erythroid parameters: red blood cells, hemoglobin, mean corpuscular volume and hematocrit. (E) Serum iron levels. (F) Liver hepcidin (*Hamp)* mRNA expression against housekeeping β-actin mRNA. (G-H) Iron content of spleen or liver. Statistical analysis was performed by two-tailed Student’s *t*-test. *\*P* < 0.05, *\*\*P* < 0.01; ns = not significant; n = 8 mice per group.

These results show that curcumin in an iron-sufficient diet causes mild anemia in a mouse model of IBD, reduces iron stores in the spleen, worsens colitis and decreases survival even in a DSS-resistant mouse strain.

## Discussion

In this study, we aimed to investigate the potential of curcumin’s chelating activity to affect body iron stores and anemia development in a murine model of IBD. We found that curcumin supplementation in DSS-treated mice led to a decrease of several erythroid parameters, including the number of red blood cells, hemoglobin, mean corpuscular volume, and hematocrit, indicating the development of mild anemia. We also showed that Curcumin-DSS mice developed splenomegaly, which is indicative of extramedullary erythropoiesis in mice responding to iron-deficiency anemia (3). These changes were accompanied by a reduction in liver hepcidin mRNA levels, further indicating that mice become anemic since hepcidin levels are inhibited by anemia (44). Hepcidin levels decreased despite the presence of inflammation, which has the opposite effect of anemia and activates hepcidin expression, confirming previous studies showing that erythropoietic drive can inhibit hepcidin activation through the inflammatory pathway (19). Our results are in agreement with previous work reporting that curcumin supplementation decreases hepcidin levels both in mice (9) and humans (29). This regulation has been associated with the inhibition of phosphorylation of the signal transducer and activator of transcription 3 (STAT3) (16, 29) and TNFα activation (48). MyD88 has also been implicated as an important factor for hepcidin regulation as MyD88-deficient mice are unable to appropriately upregulate hepcidin expression when iron-challenged and develop iron-loading in the liver (31). However, we show that curcumin does not affect MyD88 levels in the liver; hence, MyD88 would not be expected to further interfere with hepatic hepcidin expression in our experimental settings. The fact that MyD88 expression in the liver was not affected and contrasted with its downregulation in colonic tissue, indicates that curcumin regulation of MyD88 expression could be more relevant locally, within the intestine.

Our finding that curcumin modulates iron status is in line with its iron chelating properties that have been demonstrated both *in vitro* (4) and *in vivo* (22, 23). In long-term experiments in mice, curcumin supplementation has been shown to significantly lower liver and spleen iron levels (9). Others have reported that the usage of a high dosage of curcumin in a low iron diet modulates erythroid and iron parameters, exacerbating iron deficiency symptoms (23). Our present work strengthens and adds to these previous studies by demonstrating that the effect of curcumin in iron metabolism is of importance in the context of chronic intestinal inflammation.

Under our experimental conditions, Curcumin-DSS mice exhibited more severe symptoms of intestinal inflammation. This contradicts previous reports on curcumin’s anti-inflammatory effects in colitis, which had been linked to the attenuation of the TLR4/MyD88/NF-κB inflammatory pathway by inhibiting TLR4 homodimerization (58) and decreasing MyD88 expression (33). Although the reasons for these differences are not clear at present, it is worth mentioning that MyD88 deficiency in mice seems to increase susceptibility to DSS-induced colitis. Araki *et al.* reported that MyD88 is crucial in intestinal homeostasis by playing a protective role against the development of colitis since DSS induced a more severe colitis in *MyD88^-/-^* mice (1). In their experiments, they observed that mice lacking MyD88 in the colon had a subsequent higher intestinal permeability, causing more severe infiltration of bacterial products from the lumen (1). Similarly, our data show that MyD88 downregulation by curcumin in DSS-treated mice resulted in an aggravation of the inflammatory responses in our experimental conditions. In fact, DSS-curcumin mice showed more severe colonic tissue damage compared to DSS-treated mice that were not supplemented with curcumin. While the precise mechanism of action remains to be investigated, the susceptibility of MyD88-deficient mice to colitis has been linked to gut microbiota composition, which is altered in IBD (36). Furthermore, *MyD88*^-/-^ mice, in contrast with wild-type mice, are unable to respond to treatment with probiotic bacteria in the context of DSS-induced colitis, further highlighting the link between MyD88 and the gut microbiota (50). Overall, the implications of these studies are that bacterial components may play both detrimental (36) and protective roles, at least partially, in a MyD88-dependent manner (1, 50).

Regarding colitis severity, our study is in contrast with previous reports revealing a protective effect of curcumin in DSS-induced acute (13, 57) and chronic (20) colitis in mice. Such differences may be due to the amount of iron found in standard rodent chow, which tends to have excess iron ranging from 350 mg/kg up to 900 mg iron/kg diet in some related studies (20). This strikingly contrasts with our iron-sufficient diet that contained 50 mg/kg, which is more in accordance to mouse iron requirements (41). Excess dietary iron may compensate for the iron chelating effect of curcumin (2) and presumably avoids the development of iron deficiency and anemia in this model. In addition to iron, other components of the rodent diet, such as fermentable fibers, may have a role in altering the effects of curcumin in DSS-induced colitis. Indeed, recent studies have shown that the presence of fermentable fibers in the diet can ameliorate low-grade inflammation while exacerbating disease severity in response to acute colitis (38).

Previous studies have reported that the efficacy of dietary curcumin in trinitrobenzene sulfonic acid (TNBS)-induced colitis, another rodent model for IBD, may vary depending on the mouse strain (5). Furthermore, mouse strain has also been shown to influence the severity of DSS-induced colitis (37). We tested BALB/c mice since they are known to be substantially more resistant to DSS acute colitis in comparison to C57BL/6 mice (37). We found that curcumin aggravated colitis and induced an iron-deficiency anemia phenotype in BALB/c mice as well, indicating that the detrimental effect of curcumin in the context of an iron-sufficient diet is independent of mouse strain.

In conclusion, we found that long-term curcumin administration in mice has potentially adverse effects in a DSS-induced model of ulcerative colitis, lowering iron stores and leading to the development of anemia. While beneficial effects of curcumin as an anti-inflammatory agent have been documented in animal models as well as in patients with mild to moderate ulcerative colitis (30, 52), the iron chelating properties of curcumin should be taken into account. This is particularly pertinent in situations of iron-deficiency, a condition that is found in up to 78% of Crohn’s disease patients with active inflammation (43). Our study highlights the potential risks of curcumin, which is commonly taken as an over-the-counter supplement without monitoring erythroid and iron status parameters.

## Acknowledgements

This work was supported by grants from the Canadian Institutes of Health Research (CIHR, grant no. MOP123246) and the Natural Sciences and Engineering Research Council of Canada (NSERC, grant no. 412326-2011) to MMS. MC is the recipient of a postdoctoral fellowship from MITACS (Accelerate fellowship, # IT07618) and MSM received a Ph.D. scholarship from the Institute du Cancer de Montréal (ICM, bourse Canderel). We thank Jacqueline Chung for editing the manuscript.

## Author roles

MSM and MC designed and performed experiments, analyzed and interpreted data, and contributed to the writing of the manuscript. GF performed experiments and analyzed data. AC provided technical support in some experiments. MMS designed experiments, interpreted data and contributed to the writing of the manuscript. All authors read and approved the final manuscript.

